# Unravelling the constrained cell growth in engineered living materials

**DOI:** 10.1101/2025.05.21.655284

**Authors:** Shuang-Shuang Sun, Cheng-Cheng Ding, Hai-Yan Yu, Xian-Zheng Yuan, Shu-Guang Wang, Peng-Fei Xia

## Abstract

Engineered living materials (ELMs) leverage the integrative advantages of materials science and synthetic biology for advanced functionalities. Predicting and controlling cellular behavior is essential for designing and building ELMs, requiring fundamental understanding of the growth dynamics of encapsulated cells. Here, we interrogate the interference of constrained growth on the engineered functionalities and cellular physiology of cyanobacteria and unveil the dynamic interaction between cell growth and spatial confinements within photosynthetic ELMs. We observed that engineered cyanobacteria within ELMs exhibited compromised performances in growth, uptake of non-natural substrate, and synthesis of customized products, while ELMs could protect encapsulated cells from external stresses. Besides commonly accepted external influences, we identified abnormally high levels of reactive oxygen species and impaired oxygenic photosynthesis inside the cells encapsulated in ELMs. Finally, we illustrated the dynamics of cell growth within the confined spaces enveloped by the material matrices, forming clustered cell aggregates and compressed growth bubbles until the spatial limits. Our study provides a fundamental yet often overlooked connection between cellular behavior and spatial confinements, consolidating the foundation for advanced ELM innovations.

## Introduction

Engineered living materials (ELMs) are emerging field integrating synthetic biology and materials science, opening new avenues for sustainability and human health.^1–3^ Synthetic biology designs and modifies living systems to endow ELMs distinctive “living” capabilities to sense, process, and response to external environmental stimuli. Materials science, in return, facilitates the creation of ELMs by incorporating living cells within matrices.^4–6^ In addition, advanced manufacturing and processing techniques, such as 3D printing, electrospinning, and microfluidics, have been employed to fabricate ELMs with user-defined structures and morphologies, substantially expanding the potential of ELMs in environmental monitoring,^7^ biomanufacturing,^8^ diagnosis and therapeutic treatment.^9^

The functionalities of ELMs are predominantly controlled by the natural or engineered behaviors of living organisms, requesting precision prediction and control of cellular physiology. Fundamentally, cell growth within ELMs is critical for ELM innovations. For instance, photoautotrophs grow faster at the periphery of ELMs due to sufficient access to light and air, and, therefore, ELMs can be generated with increased photosynthesis via regulating the morphology.^10^ The clustered algae within ELMs exhibit shading effects, which can be alleviated by adding scattering particles for enhanced biomass accumulation.^11^ A frequently overlooked aspect is the confined environment within ELMs, where the cell growth is spatially constrained. This is a common issue in either self-organizing living materials, where the matrix is generated by the embedded living organisms themselves (e.g., biofilms and kombucha),^5,12–14^ or hybrid living materials that physically encapsulate living cells into material matrices.^15,16^ The constrained growth inevitably alters the cellular behaviors, such as the metabolic robustness and biosynthetic competency. However, the dynamic establishment and evolution of the spatial confinement along with its collateral impacts through the interactive material matrices and living cells remain unresolved, leaving a significant gap that might mislead ELM innovation efforts, hinder precision control of functionalities, and obstacle the development of biosafety measures.

Cyanobacteria are photosynthetic chassis that leverages solar energy to drive CO_2_ fixation and the metabolisms. ELMs have been designed and generated by encapsulating living cyanobacteria, e.g., *Synechococcus elongatus*, into artificial soft material matrices, such as alginate and chitosan hydrogels, showing promise for diverse applications in contaminant remediation (e.g., dyes and heavy metals),^17,18^ bioproduction,^19^ and medical treatments, including myocardial infarction treatment,^20^ post-surgery tumor prevention and wound healing.^21^ These functionalities mainly rely on the natural or engineered behaviors of cyanobacteria, for instance, the light-dependent oxygenic photosynthesis and artificial metabolic routes. In nature, *S. elongatus* grows planktonically to maximize access to light and CO_2_,^22,23^ rather than being confined within materials. This unnatural growth, though rarely explored, inevitably alters its biology and, consequently, the performance of ELMs. Therefore, *S. elongatus* not only requires but also, as a paradigm, enables deeper interrogation into the cellular behavior within ELMs.

Here, we investigated the growth dynamics of cyanobacteria within ELMs and the effects of constrained growth on functions of ELMs and cellular physiology. First, we constructed three engineered cyanobacteria, using *S. elongatus* PCC 7942 as chassis, and encapsulated these strains in alginate hydrogel, generating ELMs with distinct functionalities. Then, we systematically evaluated the performance of these functionalized ELMs in controlled cultivation conditions, comparing to the strains growing planktonically. Next, we delved into the systems-level responses of the constrained cells via biochemical and transcriptomic analysis. Finally, we interpreted the constrained cell growth within ELMs through multiple microscopic visualization approaches, establishing the evolution of spatial confinements in ELMs. Our work finds a missing puzzle piece toward a comprehensive understanding of the biology of and within ELMs, laying the foundation for future ELM innovations.

## Results

### Design of ELMs with engineered cyanobacteria

To design ELMs with engineered cyanobacteria, we chose alginate as the abiotic matrices, where alginate polymers were crosslinked with divalent cations Ca^2+^ to synergize egg-box structures.^17,24^ The resulting hydrogel allowed light to penetrate and gases (e.g., CO_2_ and O_2_) and nutrients to diffuse,^25,26^ thus sustaining the living activity of ELMs **(Figure 1A)**. We constructed three engineered *S. elongatus* PCC 7942 with distinct non-natural properties, including resistance to environmental inhibitors, expanded spectrum of substrates, and the production of value-added compounds **(Figure 1B)**. First, we integrated the *aadA* cassette, encoding the aminoglycoside-3"-adenylyltransferase, into the genome of *S. elongatus*, generating strain SS1 with spectinomycin/streptomycin resistance **(Figure 1C and Figure S1A)**. Next, we enabled glucose utilization in *S. elongatus* by complementing the glucose transporter encoded by *galP*, giving a mixotrophic strain SS2 **(Figure 1D and Figure S1B)**.^27^ Then, SS3 was constructed by inserting the sucrose permease gene *cscB* into the genome, thereby producing and secreting sucrose under salt stress **(Figure 1E and Figure S1C)**.^28^ Notably, we observed the similar phenotype of SS1 compared to the wild-type strain when cultivating without antibiotics, and we employed this strain as a control to avoid unexpected influence of selective markers in the strains **(Figure S3)**. Finally, by crosslinking the mixture of the engineered strains and alginate, we generated spherical hydrogels ELM-SS1 (with SS1), ELM-SS2 (with SS2), and ELM-SS3 (with SS3) with engineered functionalities for further evaluation.

**Figure 1.**
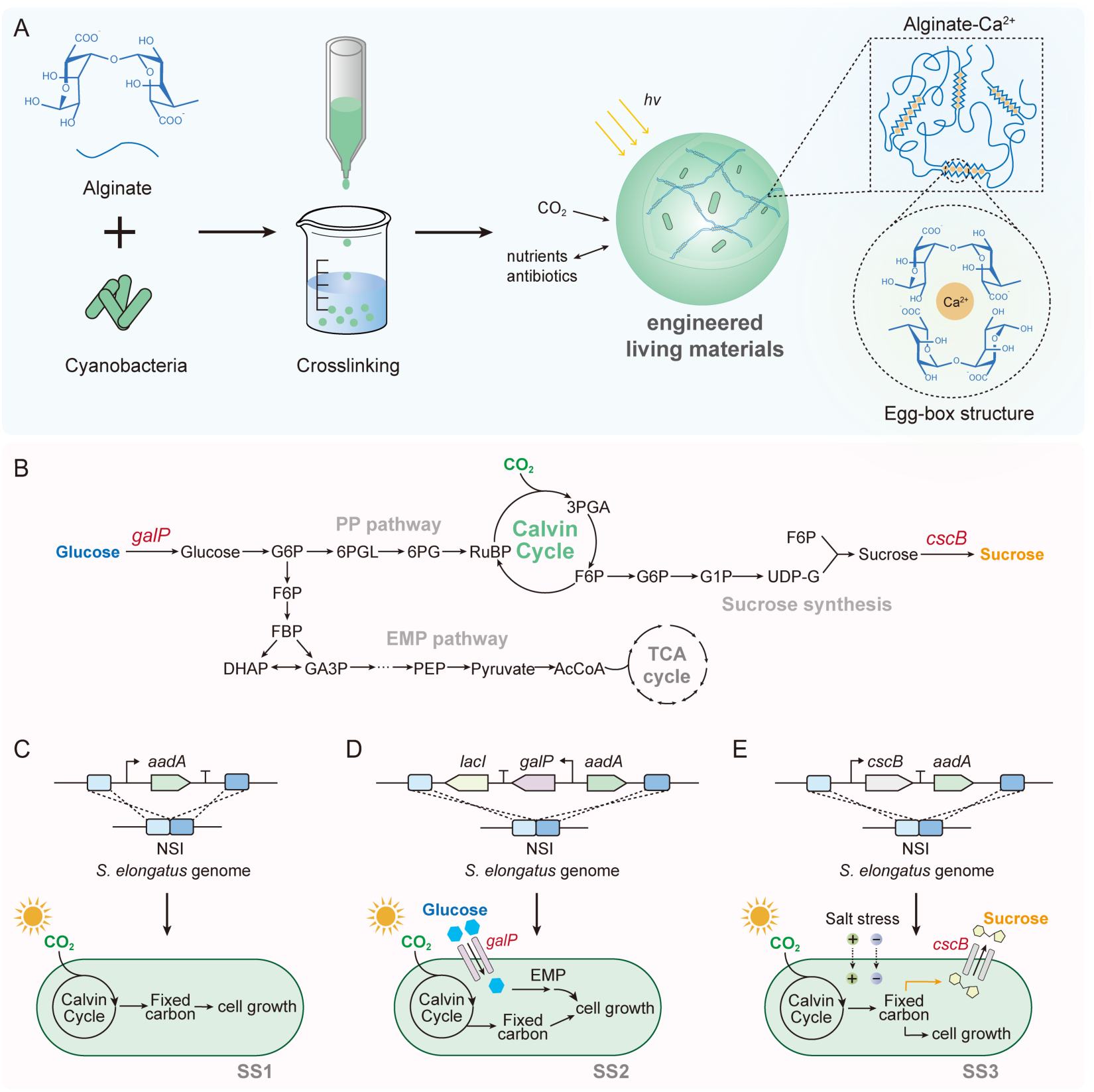
Design of photosynthetic ELMs with engineered cyanobacteria. **(A)** Schematic diagram of ELM construction. Alginate is mixed with cyanobacteria and crosslinked with CaCl_2_ to form spherical ELMs with egg-box structure, allowing light to penetrate and gases and nutrients to diffuse. **(B)** The glucose and sucrose metabolic pathway in cyanobacteria. **(C)** Construction of strain SS1 with *aadA* cassette, encoding the aminoglycoside-3"-adenylyltransferase, being integrated into the NS1 of the *S. elongatus* genome. **(D)** Construction of strain SS2 by integrating *galP*, which encodes a glucose transporter, into the NS1, allowing *S. elongatus* to utilize glucose for cell growth. **(E)** Construction of strain SS3 by inserting *cscB*, which encodes a sucrose permease, into the NS1 of the *S. elongatus* genome, leading to sucrose secretion under salt stress. PP pathway, pentose phosphate pathway; EMP pathway, glycolytic pathway; G6P, Glucose-6-phosphatase; 6PGL, 6-phosphogluconolanase; 6PG, 6-phosphogluconic acid; F6P, Fructose-6-phosphate; FBP, Fructose bisphosphate; DHAP, Dihydroxyacetone phosphate; GA3P, Glyceraldehyde-3-phosphate dehydrogenase; PEP, Phosphoenolpyruvate; AcCoA, Acetyl CoA carboxylase; RuBP, Ribulose-1,5-bisphosphate; 3PGA, 3-phosphoglycerate acid; G1P, Glucose-1-phosphatase; UDP-G, Uridine diphosphate glucose; NS1, neutral site 1.

### Compromised functionalities of ELMs compared to free-growing cyanobacteria

To evaluated the performance of ELMs, we first compared the growth of the engineered cyanobacteria within ELMs and the strains growing freely in the medium. To enable fair comparations, equal amounts of cells were utilized to generate ELMs or inoculated for cultivation, and the alginate hydrogel was dissolved before evaluating the cell growth within ELMs. We observed slower growth and lower final optical density (OD_730_) of SS1 within ELM-SS1 than growing freely in the BG11 medium **(Figure 2A and 2B)**. This result was also demonstrated by the content of chlorophyll a (Chl-a), which directly connects to oxygenic photosynthesis **(Figure 2C)**.^29^ Then, we cultivated ELM-SS2 in BG11 medium with glucose **(Figure 2A)**, and identified similarly slower growth and lower final OD_730_ of SS2 within ELM-SS2 compared to free-growing cells **(Figure 2D and 2E)**. Notably, SS2 reached a higher OD_730_ than SS1 due to the utilization of glucose **(Figure 2B and 2D)**. Within ELM-SS3, SS3 exhibited similar trends of cell growth in a medium with elevated salt stress (BG11 with 4 g/L NaCl) **(Figure 2A, 2F and 2G)**. To be noticed, we observed cell growth in the culture outside of ELMs after 8-days of cultivation **(Figure 2B-2G)**, implying cell escape from ELMs.

**Figure 2.**
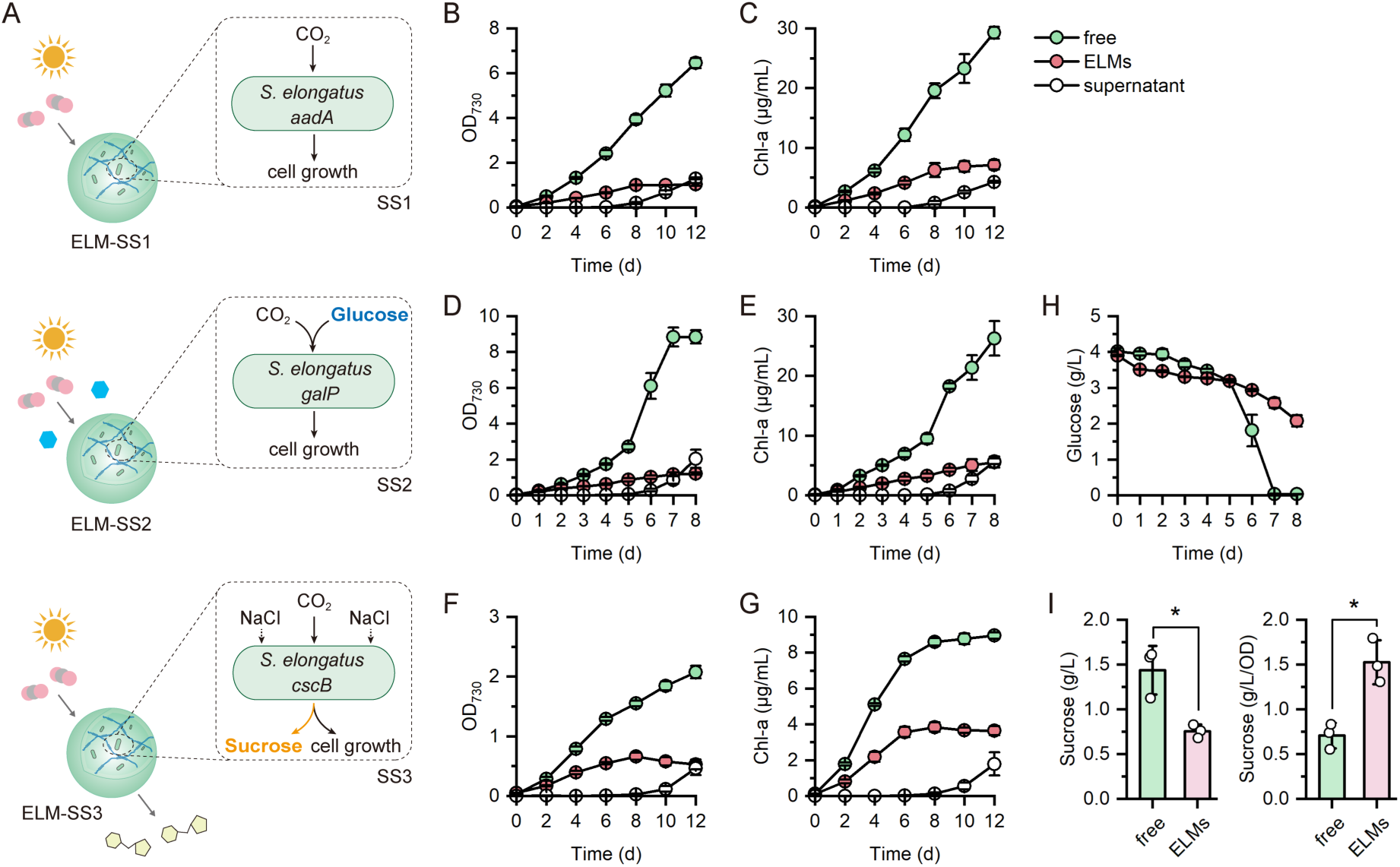
Performance of ELMs with engineered cyanobacteria. **(A)** Schematic illustration of the functionalities of the engineered ELMs. ELM-SS1 worked as control and could grow with spectinomycin/streptomycin, ELM-SS2 could fix CO_2_ and utilize glucose for cell growth, and ELM-SS3 could produce sucrose under salt stress. **(B)** OD_730_ and **(C)** the content of Chl-a of SS1 within ELM-SS1 and growing freely during the 12 days of cultivation in BG11 medium. **(D)** OD_730_ and **(E)** the content of Chl-a of SS2 within ELM-SS2 and growing freely during the 8 days of cultivation in BG11 medium with 5g/L glucose. **(F)** OD_730_ and **(G)** the content of Chl-a of SS3 within ELM-SS3 and growing freely during the 12 days of cultivation in BG11 medium with 4g/L NaCl. **(H)** Glucose consumption of ELM-SS2 and free-growing SS2. **(I)** The titer of sucrose and specific sucrose production, which was calculated by dividing the sucrose titer by OD_730_, between ELM-SS3 and free-growing SS3 on Day 12. All experiments were performed in triplicate, and the error bars indicate the standard deviations of the means of biological replicates. The differences were statistically assessed by the *t*-test (\**P* < 0.05).

Subsequently, we evaluated the functionalities of ELM-SS2 and ELM-SS3. As designed, ELM-SS2 could use glucose as an additional carbon source for cell growth, while the consumption of glucose by ELM-SS2 was slower than that of SS2 growing freely in the medium **(Figure 2H)**. Moreover, we detected sucrose production of ELM-SS3 in BG11 medium with 4 g/L NaCl, and the titer of sucrose was lower than that of SS3 alone **(Figure 2I and Figure S4)**. However, when we calculated the specific sucrose production through dividing sucrose production by OD_730_, SS3 in ELM-SS3 showed superior performance than free-growing SS3 **(Figure 2I)**, which were in agreement with another report.^19^ Two reasons might attribute to the repressed growth of encapsulated cells and impeded performance of ELMs. One is the material matrices slow down the diffusion of CO_2_, nutrients, or other substrates (e.g., glucose),^10^ while the other is the self-shading effect between the ELMs limiting accessible light.^30^ The superior performance of SS3, on the other hand, suggesting distinct variations in cellular physiology within ELMs.

### ELMs protect cyanobacteria from environmental fluctuations

We further investigated the performance of ELMs under environmental fluctuations. Light is the most critical factor influencing the growth of cyanobacteria.^31^ It has been reported that strong illumination represses photosystem II (PSII), known as photoinhibition.^32^ Therefore, we chose light intensity as the primary environmental fluctuation and cultivated ELM-SS1 and free-growing cells under strong (70 μmol photons⋅m^-2^⋅s^-1^) and normal light (35 μmol photons⋅m^-2^⋅s^-1^) **(Figure 3A)**. We found that the cell growth of SS1, demonstrated by both the OD_730_ (**Figure 3B**) and Chl-a (**Figure 3C**), were severely inhibited when growing freely in the medium under strong light. The differences were certified via statistical analysis **(Figure 3D and 3E)**. To the contrary, SS1 grew similarly within ELM-SS1 under these two light intensities **(Figure 3B and 3C)**, and the final OD_730_ and Chl-a content were at an equivalent level. While the differences of the Chl-a content in SS1 within ELMs were significantly different due to the strong illuminations, the decreases were marginal in absolute values **(Figure 3D and 3E)**. To further validate the phenomenon, we cultivated ELM-SS2 and ELM-SS3 under the two different light intensities and compared them with strains growing freely. As expected, similar growth profiles were identified **(Figure S5 and S6)**, suggesting a universal photoprotective effect of ELMs on the engineered cyanobacteria.

**Figure 3.**
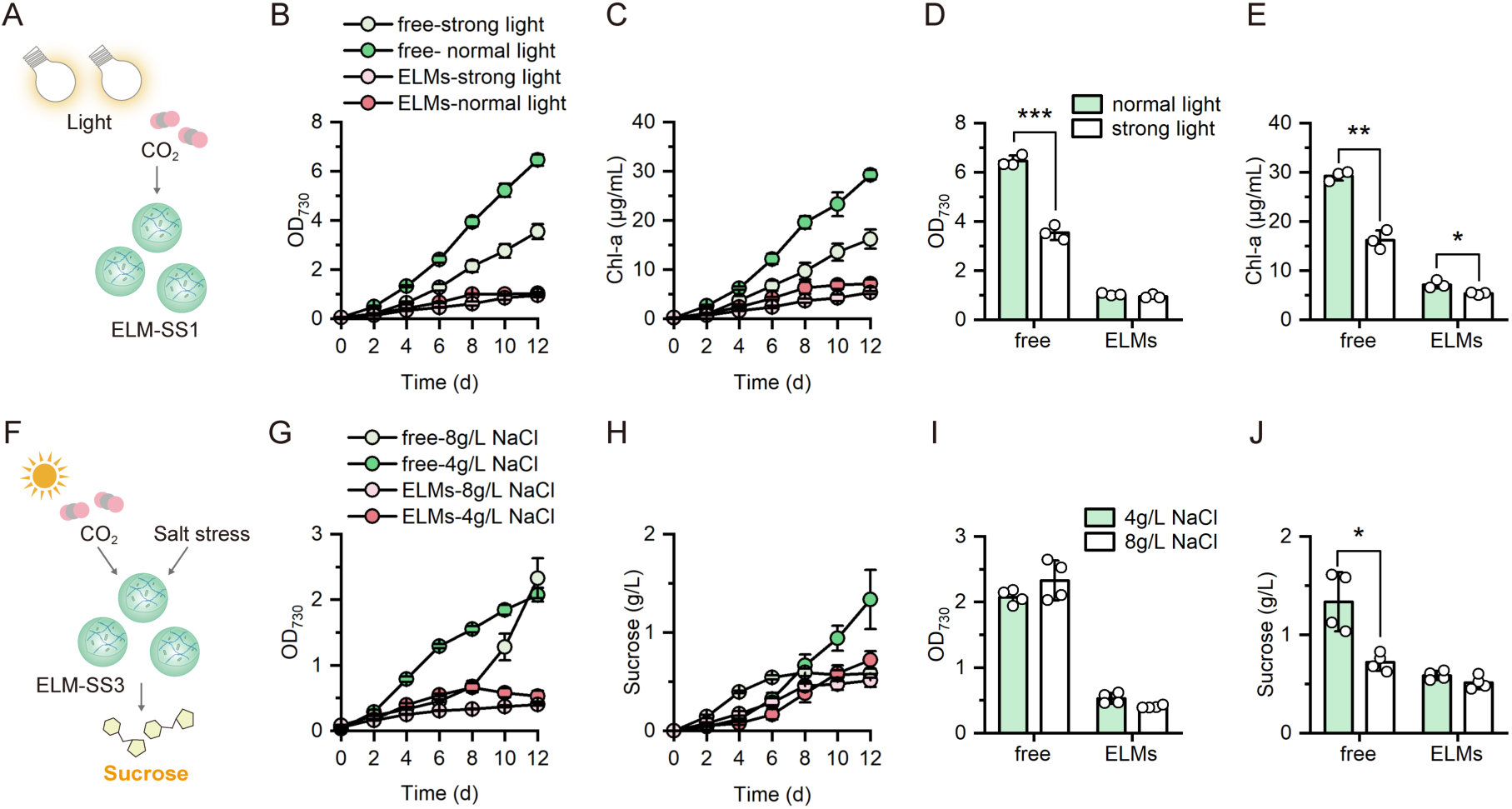
Protective role of ELMs against environmental fluctuations. **(A)** Illustration of ELM-SS1 cultivation with strong illumination as environmental fluctuation. **(B)** OD_730_ and **(C)** the content of Chl-a of SS1 within ELM-SS1 and free-growing SS1 during the 12 days of cultivation at different light intensities. **(D)** OD_730_ and **(E)** the content of Chl-a of SS1 in ELM-SS1 and growing freely on Day 12 at different light intensities, where 70 μmol photons⋅m^-2^⋅s^-1^ was regarded as strong light intensity, and 35 μmol photons⋅m^-2^⋅s^-1^ was considered normal light intensity. **(F)** Schematic diagram of sucrose producing ELM-SS3 under salt stress. **(G)** OD_730_ of SS3 within ELM-SS3 and free-growing SS3 under different salt stresses (4 g/L and 8 g/L NaCl). **(H)** The titer of sucrose secreted by ELM-SS3 and free-growing SS3 under different salt stresses. **(I)** OD_730_ of SS3 in ELM-SS3 and free-growing SS3 on Day 12. **(J)** Sucrose production of ELM-SS3 and free-growing SS3 on Day 12. The experiments were performed at least in triplicate, the error bars indicate the standard deviations of the means of biological replicates, and the differences were determined by the *t*-test (\**P* < 0.05; \*\**P* < 0.01; \*\*\**P* < 0.001).

We selected salt stress as another environmental fluctuation, which is also an essential factor for sucrose production, to evaluate the performance of ELMs **(Figure 3F)**. Increased salt stress has been reported to disrupt intracellular ion homeostasis, and, thereby, impair cellular physiologies.^33^ We observed that the growth of free-growing SS3 was severely inhibited and the sucrose production was promoted with an increased salt stress (8 g/L NaCl) at the first 6 days **(Figure 3G and 3H)**. But, the growth was rescued since Day 6 and finally reached a comparable OD_730_ to that grown in 4 g/L NaCl **(Figure 3G and 3I)**. At the same time, the sucrose production terminated, indicating a substantial alteration in cellular physiology induced by high salt stress. The growth of SS3 in ELM-SS3 was also inhibited in high salt stress, while no sudden changes in growth were identified **(Figure 3G)**. Under different salt stress, ELM-SS3 exhibited comparable performance in sucrose production during the 12 days of cultivation **(Figure 3H and 3J)** with a similar specific sucrose production as well **(Figure S7)**. These results demonstrated that ELMs can alleviate the adverse impacts of external disturbance on the biological functionalities of engineered cyanobacteria.

### Unusual oxidative stress and transcriptional responses

Besides the commonly accepted influences of mass transfer and light accessibility,^10,11^ constrained cell growth inevitably alters the cellular behavior, exerting a key challenging in precision predicting and programming biological systems. To reveal the impacts of constrained growth, we evaluated the possible oxidative stress and corresponding responses, which is a common consequence under sub-optimal conditions, via detection of reactive oxygen species (ROS), glutathione (GSH) content, and superoxide dismutase (SOD) activities.^34^ Taking ELM-SS1 as an example, we found a surprisingly high level of intracellular ROS in SS1 within ELM-SS1, which was 4.58-fold higher than that in free-growing cells **(Figure 4A)**, while no significant differences in the antioxidant GSH content and SOD activity were observed **(Figure 4B and 4C)**. In addition, two important redox equivalents, NADH and NADPH, were also determined, and we found that NADH/NAD^+^ and NADPH/NADP^+^ were significantly increased in SS1 of ELM-SS1 **(Figure 4D and 4E)**. These results implied unusual intracellular physiologies of the constrained cells, which might have collateral influence back on cell growth and functions.

**Figure 4.**
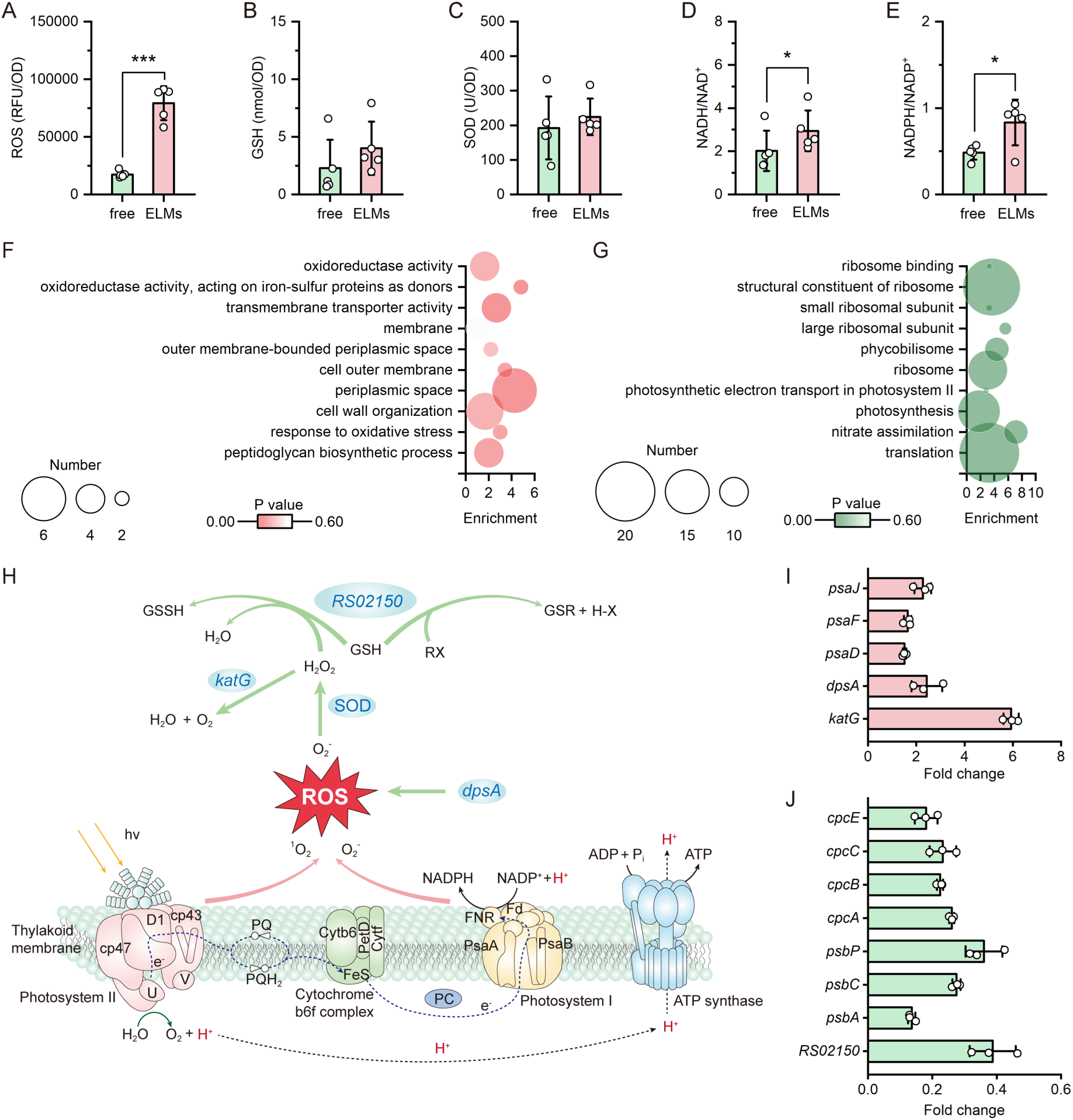
Systems-level variations induced by confined growth. **(A)** Intracellular ROS levels, **(B)** GSH content, **(C)** SOD activity, **(D)** NADH/NAD^+^, and **(E)** NADPH/NADP^+^. Green color represents free-growing SS1, while red color represents SS1 within ELM-SS1. **(F)** Representative upregulated GO terms of DEGs, and **(G)** representative downregulated GO terms of DEGs. The color indicates P values, and the bubble size indicates the number of DEGs in GO terms. **(H)** Schematic illustration of the responses related to oxidative stress and oxygenic photosynthesis in *S. elongatus* growing in constrained spaces. Fold change of **(I)** upregulated expressed genes and **(J)** downregulated expressed genes in the schematic diagram. The experiments were performed at least in triplicate, the error bars indicate the standard deviations of the means of biological replicates. The differences were statistically evaluated by *t*-test (\**P* < 0.05; \*\*\**P* < 0.001).

We performed RNA sequencing and analyzed the global responses of constrained cells at the transcription level. A total of 560 differentially expressed genes (DEGs) were identified by comparing SS1 within ELM-SS1 with SS1 growing freely **(Figure S8)**. We found that constrained growth may affect the transcription, translation and amino acid metabolism in *S. elongatus*. Using the Gene Ontology (GO) database to identify the functional categories of DEGs, we found that multiple upregulated GO terms were related to oxidative stress **(Figure 4F and Table S5)**, which was consistent with the biochemical analysis above. We further utilized the Kyoto Encyclopedia of Genes and Genomes (KEGG) database to analyze the responses of oxidative stress within ELMs **(Figure S9 and Table S6)**. Particularly, the *katG* gene, encoding the hydroperoxidase that converts H_2_O_2_ to H_2_O and O_2_,^35^ was significantly upregulated **(Figure 4H and 4I)**. In addition, the *dpsA* gene was significantly upregulated **(Figure 4I)**, *dpsA* encodes a DNA-binding hemoprotein that protects cells from oxidative stress by scavenging ROS **(Figure 4H)**.^36^ Conversely, *RS02150* gene, encoding the glutathione S-transferase (GST), was downregulated **(Figure 4J)**. GST plays a key role in the decomposition of H_2_O_2_ and the detoxification of harmful compounds through catalyzing the reduced chemical GSH into GSSH and glutathione conjugates (GSR), respectively **(Figure 4H)**.^37^ These results suggested that the constrained cells struggled to generate antioxidants to prevent oxidative damage, though the escalated ROS level exceeded the capacity.

Notably, we observed downregulated GO terms involved in ribosome and photosynthesis **(Figure 4G)**. The ribosome is the place where mRNAs are decoded and translated to peptides, forming functional proteins subsequently inside a living cell.^38,39^ The impairment of ribosome might be one reason leading to repressed cell growth. Cyanobacteria are photoautotrophic microorganisms relying on protein complexes embedded in thylakoid membranes to carry out photosynthesis, including photosystem I (PSI), PSII, cytochrome b6f complex (cytb6f), and ATP synthase. These complexes facilitate the conversion of light energy into chemical energy, producing ATP and NADPH and providing energy for carbon fixation **(Figure 4H)**.^40,41^ We found that *psbA* gene was downregulated **(Figure 4J)**, which encodes the photoreaction center protein D1 in PSII **(Figure 4H)**.^42^ Besides, oxygen evolving complex (OEC) (encoded by *psbP*) and antenna proteins (encoded by *pcbC* and *cpc* gene clusters) were downregulated **(Figure 4J)**. Conversely, the genes related to PSI were upregulated, including *psaD, psaF* and *psaJ* **(Figure 4I)**. The up-regulated PSI might contribute to the increased NADPH to NADP^+^ ratio and lead to a decrease in the ratio of PSII to PSI,^43^ which ultimately repressed the oxygenic photosynthesis of constrained *S. elongatus*.

### Cyanobacteria are constrained in growth bubbles within ELMs

The functionalities and cellular physiology were closely correlated to the growth of encapsulated cells, while the exact growth dynamics and the evolution of spatial confinement remain unclear. To decipher the cell growth within ELMs, we chose ELM-SS1 as the representative. Visualized by scanning electron microscopy (SEM), we identified a lamellar porous structure of ELM-SS1 during cultivation **(Figure 5A and Figure S11A)**, and the individual ELM bead turned from light green to dark during 12 days of cultivation due to cell growth **(Figure 5B)**. We first analyzed the surface morphology and the mechanical properties of ELMs using atomic force microscopy (AFM). The results showed increased number of small particles on the surface of ELMs along with cultivation **(Figure 5C and Figure S11B)**. The surface roughness increased sharply at the first 4 days, and, then, decreased from Day 4 to Day 12 with the overall roughness higher than Day 0. The decreased roughness might due to the evenly distributed particles after 4 days **(Figure 5C and 5D)**. Notably, the Young’s modulus exhibited a similar trend to roughness while the mechanical strength dropped significantly at Day 12 **(Figure 5E)**. In agreement with AFM analysis, we identified small bubble-like bulges on the surface of ELMs using SEM **(Figure 5F and Figure S11C)**.

**Figure 5.**
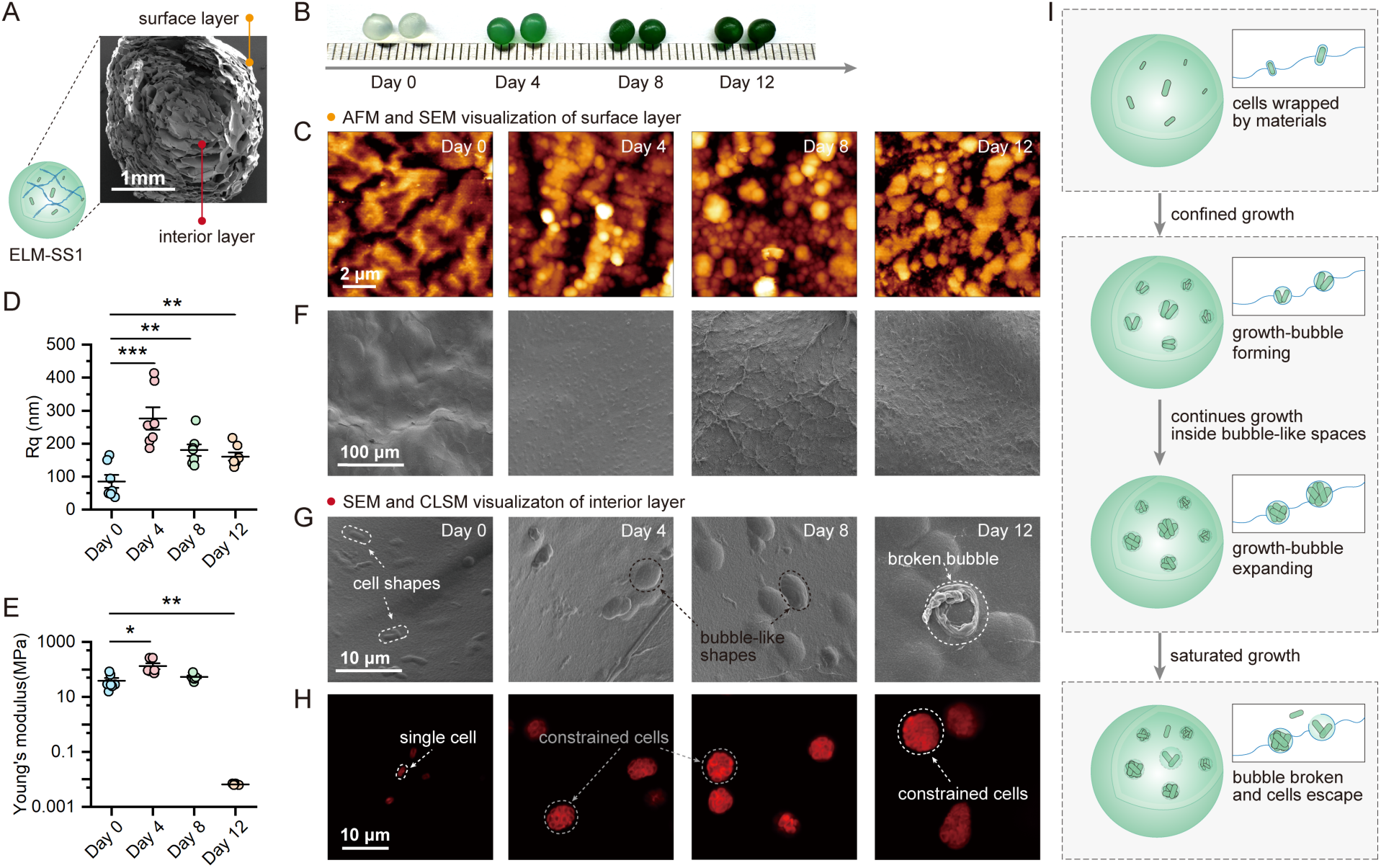
Growth dynamics of confined cyanobacteria within ELMs. **(A)** The general structure of ELM-SS1 visualized by SEM. **(B)** Color variation of ELM-SS1 beads along with cultivation. **(C)** Surface morphology of ELM-SS1 visualized by AFM. **(D)** The surface roughness and **(E)** the Young’s modulus of ELM-SS1. The differences were statistically determined by the *t*-test (\**P* < 0.05; \*\**P* < 0.01; \*\*\**P* < 0.001). **(F)** Surface morphology and **(G)** interior structure of ELM-SS1 visualized by SEM. **(H**) Representative CLSM images of ELM-SS1. Red fluorescence represents chlorophyll autofluorescence of *S. elongatus* cells. (**I**) Schematic diagram illustrating the dynamics of confined growth of *S. elongatus* cells within ELMs. All analysis were performed on samples at 0, 4, 8 and 12 days of cultivation.

Next, we analyzed the interior of ELMs to unveil the growth of encapsulated cells. Interestingly, we could not identify *S. elongatus* cells inside the ELMs. Instead, we identified cell-shaped bulges at Day 0 and bubble-like structures dispersed on the lamellae within ELMs since Day 4, which expanded along with cultivation **(Figure 5G and Figure S11D)**. These bubble-like structures were also documented in another report.^18^ We hypothesized that cyanobacteria were first wrapped with alginate polymers, forming the cell shaped bulges, and the growth of cells was constrained to these limited spaces, where the newly divided cells could only live inside and struggle to expand the space, eventually forming the observed growth bubbles. To demonstrate the hypothesis, we utilized the confocal laser scanning microscope (CLSM) to visualize the *S. elongatus* cells within ELMs based on the chlorophyll autofluorescence. If the growth of *S. elongatus* was indeed confined in these bubbles, spherically aggregated red fluorescence would be identified rather than dispersed cell shaped fluorescence. As expected, we observed cell-shaped fluorescence at Day 0, and along with cell growth, we found the fluorescence clustered together and expanded, agreeing with the SEM visualizations and demonstrating the confined cellular activities in these growth bubbles **(Figure 5H and Figure S11E)**. Furthermore, we identified the break of growth bubbles and cell escape at Day 12 **(Figure 5G)**. This could explain the escaped cells in the culture in the previous experiments **(Figure 2B-2G)**, providing the first and direct evidence on the mechanisms of cell escape from ELMs. Notably, we observed the cell morphologies were mechanically compressed in these growth bubbles, changing from a rod shape to ellipsoidal **(Figure 5H)**, which might be a direct consequence of confined growth and eventually lead to the abnormally high-level of ROS and varied physiology.^44^

To illustrate the dynamic interactions, the engineered cells were wrapped by the material matrices and confined in the resulting space when generating the ELMs. Then, the cells can only grow and being compressed in these constrained spaces forming growth bubbles. The bubbles were expanded due to cell growth, while, in turn, exerted mechanical compression leading to physiological variations. When the growth bubble reached the maximal capacity, it would break and release the cells, eventually completing the life-time dynamics of the confined growth within ELMs **(Figure 5I)**.

## Discussion

Predicting and controlling the cell behavior is critical in designing and building ELMs. Here, we unravel the confined growth of engineered cyanobacteria within ELMs and the collateral influences on ELMs functionalities and cell physiology. We found that engineered *S. elongatus* PCC 7942, which is a naturally planktonic cyanobacteria, exhibited compromised performances in growth, uptake of non-natural substrate, and synthesis of customized products within ELMs **(Figure 2)**, while ELMs protected the strains from external stresses **(Figure 3)**. These fundamental but often overlooked comparisons between encapsulated and free-growing cells provide important guidance for ELM innovations. For instance, ELMs might not be an ideal candidate for biorefinery and bioproduction where maximal productivity is demanded, while they can show stable performance under highly fluctuating conditions. Moreover, the turn-over of ELMs requires not only cell growth but also regeneration of material matrices.

Photosynthetic ELMs are not always seeking superior CO_2_ fixation and photosynthesis but also unique advantages merging materials science and synthetic biology, making precision prediction of cellular behaviors even more important. We found that the confined growth led to abnormally high level of ROS and significantly impaired oxygenic photosynthesis. These variations can be one of the reasons responsible for the repressed growth. Besides, ROS lies in the center of multiple cellular behaviors. As a consequence of confined growth, ROS can induce diverse physiological changes, including ribosome impairment, affecting signaling, and metabolic regulation.^45–47^ From a distinct perspective, the escalated ROS levels might also open new avenues for ELM designs, for instance, degradation of organic compounds ^48,49^ and induction of host immunity.^50^ Medical applications of ELMs rely on the light-dependent oxygenic activities, the downregulated PSII and OEC should be counted for fine regulation of these process, thereby achieving optimized performances.

These phenotypical and physiological variations are not only attributed to the diminished access to substrate, nutrients and light but also the confined growth of cells that are compressed and struggling to grow in limited spaces. We discovered that the engineered cells were enveloped by the alginate polymers when synthesizing the ELMs. The cell could only grow inside these confined spaces, forming clustered cell aggregates and growth bubbles dispersed on the lamellae within ELMs. Along with cultivation, the grown cells expanded the growth bubbles till their capacity limits, while, as feedback, were compressed. This suggests a previously missed connection between varied cell behavior and the spatial confinement through mechanical compression. Compression-induced physiological changes, such as escalated ROS, have been well documented in mammalian cells and human cell lines,^44^ while this is the first report illustrating such mechanism in ELMs. Eventually, the bubbles would break due to the physical properties of the material and release the cells inside **(Figure 5I)**. This encapsulation is different from self-organized ELMs, such as biofilms and kombucha, where the extracellular matrix and cells mixed together and the cells are not actually physically contained.^51,52^

While the aggregated cell growth within ELMs have been noticed for microalgae,^10,11^ our observations, for the first time, integrates the abiotic materials and living biosystems to establish the dynamics of cell growth and evolution of spatial confinements. An immediate influence of our finding is providing evidenced explanation of cell escapes from ELMs. Understanding the escape of cells is essential in design more stable ELMs, and it also lays the foundation for assessing biosafety issues and developing corresponding containment strategies.^24,53,54^ Taking together, our observations provide an indispensable building block to lay the foundation of ELM innovations, and navigate ELMs find their ideal niche to promote sustainability and human health.

## Methods

### Strains and cultivation conditions

All strains used in this study are listed in **Table S1**. *Escherichia coli* DH5α (Takara Biotech.) for molecular cloning was cultivated at 37 °C in Luria Bertani (LB) medium, consisting of 10 g/L tryptone, 5 g/L yeast extract, and 10 g/L NaCl. Agar (1.5%) was added for solid medium, and spectinomycin (60 µg/mL) was added for selection and maintaining plasmids in *E. coli*. *S. elongatus* PCC 7942 was cultivated in BG11 medium with continuous illumination (35 µmol protons⋅m^-2^×s^-1^) at 30 °C and 100 rpm. Strain SS1, SS2 and SS3 were cultivated in BG11 medium containing spectinomycin (2 µg/mL) and streptomycin (2 µg/mL) with continuous illumination. In addition, 5 g/L glucose and 0.1 mM IPTG were added for SS2 cultivations, strain SS3 was cultivated with 4 g/L or 8 g/L NaCl, and two light intensities were utilized to evaluate the effects of illumination, with 35 µmol protons⋅m^-2^×s^-1^ as normal light intensity and 70 µmol protons⋅m^-2^×s^-1^ as strong light intensity.

### Plasmid construction and transformation of cyanobacteria

All plasmids used in this study are listed in **Table S1**, and the primers are listed in **Table S2**. The plasmid pAM2991 was used for gene integration into the NS1 in the *S. elongatus* genome. Plasmid pAM2991-Δ*lacI* was constructed via knocking out *lacI* cassette (**Figure S1A**). pAM2991-Δ*lacI*-*cscB* was generated by inserting *cscB* cassette into pAM2991-

Δ*lacI* via DNA assembly (**Figure S1C**). PCR was performed using PrimeSTAR Max DNA

Polymerase (Takara Biotech.) and DNA assembly was conducted using In-Fusion Snap

Assembly Premix Kit (Takara Biotech.). All plasmids were extracted using TIANprep Mini Plasmid Kit (Tiangen Biotech.), verified by gel electrophoresis (**Figure S2**) and quantified by NanoDrop One (Thermo Fisher Scientific). The *cscB* cassette was codon optimized and synthesized by Beijing Liuhe BGI, and the sequence of *cscB* cassette can be found in **Table S3**.

To transform cyanobacteria, *S. elongatus* PCC 7942 was cultivated until OD_730_ reaching 0.5-0.7, and then 15 mL of culture was collected and centrifuged at 5,000 rpm for 15 min. The cells were washed by BG11 and resuspended in 300 µL BG11, 2 µg of plasmid was added, and the cells were incubated at 30 °C in the dark for 16-18 h. Subsequently, the cells were cultured with BG11 containing antibiotics in light for 24 h at 30 °C. Finally, the cells were plated on BG11 plates with antibiotics for selection, and then incubated statically in light at 30 °C. Spectinomycin (2 µg/mL) and streptomycin (2 µg/mL) were added for selection of transformants.

### Colony PCR and Sanger sequencing

After transformation, single colonies were randomly selected for PCR. First, the colony was resuspended in 20 µL of ddH_2_O and lysed at 100 °C for 10 min. Next, 1 µL of supernatant was utilized as the template DNA, 0.5 µL of each primer, 10 µL of the PrimeSTAR Max DNA Polymerase master mix (Takara Biotech.) and 8 µL ddH_2_O were mixed for PCR. The PCR products were determined via Sanger sequencing. The primers used for colony PCR and Sanger sequencing are listed in **Table S2**.

### Preparation and dissolution of ELMs

All engineered cyanobacteria were cultivated in BG11 until OD_730_ 0.5-0.7 for preparing the living component of ELMs. 5% (w/v) alginate solution was prepared and sterilized at 121 °C for 25 min. The above engineered cyanobacteria were mixed with sterile alginate solution in a 1:1 (v/v) ratio to reach a final alginate concentration of 2.5% (w/v). Then, the mixture of cyanobacteria and alginate was transferred into a 10 mL sterile syringe, where the mixture was squeezed out and crosslinked by dropping it into 5% (w/v) sterilized CaCl_2_ solution to form ELMs. Finally, the CaCl_2_ solution was removed, and ELMs were washed twice with sterile water. The ELMs were dissolved in 3% (w/v) sodium citrate solution when necessary. The mixture was centrifuged at 15,000 rpm for 10 min, the supernatant was discarded and the cell pellets were resuspended to their original sample volumes using fresh BG11 medium for subsequent assays.

### Determination of cell growth, glucose and sucrose

The growth was characterized by OD_730_ and the content of Chl-a. Chl-a analysis was following a reported protocol.^55^ Briefly, 1 mL of cell was centrifuged at 15,000 rpm for 10 min, and, after discarding the supernatant, the cells were resuspended in 1 mL precooled methanol and held at 4 °C in the dark until the green color was disappeared in the cell pellet. The absorbance of supernatant was measured at 665 nm and 720 nm using UV-vis spectrophotometer (T600, PERSEE), and the content of Chl-a was calculated by the following equation:

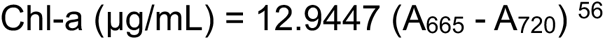

Glucose was detected using HPLC (Agilent 1260) with a reflectance index detector (RID) at 50 °C. The mobile phase was sulfuric acid (0.5 mM) at a flow rate of 0.6 mL/min. Sucrose was measured using plant sucrose content assay kit (Solarbio, China) following the manufacturer’s protocol. The specific sucrose production was calculated via dividing sucrose production by OD_730_, representing the capacity of sucrose production of individual cells.

### Microscopical characterization of ELMs

ELM-SS1 was collected and washed twice with fresh BG11, and then immobilized on petri dish using Pattex AB glue for 10 min. Fresh BG11 was added until ELM-SS1 was fully immersed. AFM (JPK Nano-Wizard 4, Bruker, Germany) analysis was performed using a quadratic pyramid probe, and the surface roughness and the Young’s modulus were analyzed by JPKSPM Date Processing software. For SEM (Quanta 250 FEG, FEI, USA) sample preparation, the washed ELM-SS1 were frozen in liquid nitrogen for 5 min and dried through freeze vacuum drying method. The lyophilized samples were adhered to SEM sample stubs using double-sided carbon tape and coated with gold particles for 200 sec. The washed ELM-SS1 was visualized by CLSM (LSM 880, Zeiss, Germany) with excitation wavelength of 561 nm for chlorophyll-autofluorescence.

### Analysis of oxidative stress

ROS was determined by a DCFH-DA fluorescent probe kit (Nanjing Jiancheng, China), and the fluorescence was detected at 488 nm excitation and 525 nm emission wavelength. All samples were lysed using an ultrasonic crusher. GSH and SOD were determined using specific assay kits (Nanjing Jiancheng, China), and the absorbance was measured at 405 nm and 450 nm, respectively. NADH/NAD^+^ and NADPH/NADP^+^ were determined using specific assay kits (Beyotime Biotechnology, China), and the absorbance was measured at 450 nm. All measurements were performed using a multimode microplate reader (Spark, Tecan) and the concentrations were normalized to OD_730_ before statistical analysis. All oxidative stress indictors are listed in **Table S4**.

### Transcriptomics and quantitative PCR (qPCR) analysis

ELM-SS1 and free-growing cells were cultivated in BG11 medium until exponential phase (OD_730_ 0.5-0.7). First, ELM-SS1 were collected and fully dissolved. Then, all samples were washed with precooled phosphate buffered saline (PBS) for three times, frozen in liquid nitrogen for 5 min and stored at −80 °C until analysis. The RNA sequencing (RNA-seq) data have been stored in the National Center for Biotechnology Information (NCBI) SRA database under accession number PRJNA1240180. The *katG* and *RS02150* were selected to validate the differentially expressed genes (DEGs) by quantitative PCR (qPCR), which was performed using Applied Biosystems QuantStudio 5 (Thermo Fisher Scientific, USA) (**Figure S10)**. The gene encoding 16S ribosomal RNA was used as the housekeeping gene. All primers used for qPCR are listed in **Table S2**.

### Statistical analysis

All experiments were performed at least in triplicate, and the error bars indicate the standard deviations of the means of biological replicates. The differences were statistically determined via the *t*-test (\**P* < 0.05; \*\**P* < 0.01; \*\*\**P* < 0.001).

## Conflict of Interests

The authors declare no conflict of interest.

## Acknowledgments

This work was supported by the National Natural Science Foundation of China (22278246 and 22378233), the Department of Science and Technology of Shandong Province (2022HWYQ-017), the Qilu Young Scholar Program of Shandong University (to P.-F.X.), the Taishan Scholars Project of Shandong Province (NO. tstp20230604), and the Intramural Joint Program Fund of State Key Laboratory of Microbial Technology (Project NO. SKLMTIJP-2024-01). The authors thank the support from Analytical Testing Center in School of Environmental Science and Engineering at Shandong University.

## Notes

### Competing Interest Statement

The authors have declared no competing interest.

