## Supporting Information for "Unravelling the constrained cell growth in engineered living materials"

**Table S1.** Strains and plasmids used in this study

| | **Description** | **Source** |
|---|---|---|
| **Strains** | | |
| *E. coli* DH5α | Commercial *E. coli* strain for molecular cloning | Takara Biotech. |
| *S. elongatus* PCC 7942 | Wild-type strain | Lab stock (ATCC 33912) |
| SS1 | PCC 7942, *aadA* cassette integrated into NS1 | This study |
| SS2 | PCC 7942, *galP* gene integrated into NS1 | Lab stock |
| SS3 | PCC 7942, *cscB* gene integrated into NS1 | This study |
| **Plasmids** | | |
| pAM2991 | *colE1* ori, *lacI*-$P_{trc}$, *aadA* cassette | Lab stock (Addgene #40248) |
| pAM2991-Δ*lacI* | *colE1* ori, *aadA* cassette | This study |
| pAM2991-Δ*lacI-cscB* | *colE1* ori, *aadA* cassette, *cscB* | This study |

**Table S2.** Primers used in this study

| Primer | Sequence | Target gene |
|---|---|---|
| **Primers for Inverse PCR** | | |
| XIA-SS-003 | AGATCGCTAACTGGTTGACGTAAATGCATGCCGCTTC | |
| XIA-SS-004 | CAACCAGTTAGCGATCTGTTAGCGCGAATTGATCTGG | |
| **Primers for In-Fusion assembly** | | |
| XIA-SS-016 | GCCGCCATGAGCGGATACATTTGACGGCTAGCTCAGTCCT | |
| XIA-SS-017 | GACATCATAACGGTTCTGGCATTTGTCCTACTCAGGAGAG | |
| XIA-SS-018 | CTCTCCTGAGTAGGACAAATGCCAGAACCGTTATGATGTC | |
| XIA-SS-019 | AGGACTGAGCTAGCCGTCAAATGTATCCGCTCATGGCGGC | |
| **Primers for colony PCR and Sanger sequencing** | | |
| XIA-SS-022 | CGATCAACTTGCGAGCAGCC | |
| XIA-SS-023 | CGGCGTCGGCTTGAACGAAT | |
| XIA-SS-025 | CCACGTCGATATCTGGCACG | |
| XIA-SS-027 | TTGCCAAGGAGACCCTCGAA | |
| XIA-SS-028 | TATCGGCGCTTTCTTCGCTG | |
| XIA-SS-029 | ATGCGGGACAACGTAAGCAC | |
| XIA-SS-030 | GGCGGTTTTCATGGCTTGTT | |
| **Primers for qPCR** | | |
| XIA-SS-048 | TCTGAGAGGATGATCAGCCA | 16S rRNA |
| XIA-SS-049 | TTCTTCCCTGAGAAAAGGGG | 16S rRNA |
| XIA-SS-050 | TGCCGTCCACTTCACCTGAT | *katG* |
| XIA-SS-051 | ACGAAGCACGGTGTCTTCAC | *katG* |
| XIA-SS-052 | AGCGCCGATCGCAAGCACAG | *RS02150* |
| XIA-SS-053 | CTGGTTGTTTGCAGGCTTCG | *RS02150* |

**Table S3.** The sequence of *cscB* cassette.

| **Sequence** |
|---|
| TTGACGGCTAGCTCAGTCCTAGGTACAGTGCTAGCAATTTCACACAGGAAACAGACCATGGCTCTG |
| AACATCCCCTTCCGCAACGCTTACTACCGCTTCGCTAGCAGCTACAGCTTCCTGTTCTTCATCAGC |
| TGGAGCCTGTGGTGGAGCCTGTACGCTATCTGGCTGAAAGGCCACCTGGGCCTGACCGGCACCG |
| AACTGGGCACCCTGTACAGCGTGAACCAATTCACCAGCATCCTGTTCATGATGTTCTACGGCATCG |
| TGCAAGACAAACTGGGCCTGAAAAAACCCCTGATCTGGTGCATGAGCTTCATCCTGGTGCTGACC |
| GGCCCCTTCATGATCTACGTGTACGAACCCCTGCTGCAAAGCAACTTCAGCGTGGGCCTGATCCT |
| GGGCGCTCTGTTCTTCGGCCTGGGCTACCTGGCTGGCTGCGGCCTGCTGGACAGCTTCACCGAA |
| AAAATGGCTCGCAACTTCCACTTCGAATACGGCACCGCTCGCGCTTGGGGCAGCTTCGGCTACGC |
| TATCGGCGCTTTCTTCGCTGGCATCTTCTTCAGCATCAGCCCCCACATCAACTTCTGGCTGGTGAG |
| CCTGTTCGGCGCTGTGTTCATGATGATCAACATGCGCTTCAAAGACAAAGACCACCAATGCGTGG |
| CTGCTGACGCTGGCGGCGTGAAAAAAGAAGACTTCATCGCTGTGTTCAAAGACCGCAACTTCTGG |
| GTGTTCGTGATCTTCATCGTGGGCACCTGGAGCTTCTACAACATCTTCGACCAACAACTGTTCCCC |
| GTGTTCTACAGCGGCCTGTTCGAAAGCCACGACGTGGGCACCCGCCTGTACGGCTACCTGAACA |
| GCTTCCAAGTGGTGCTGGAAGCTCTGTGCATGGCTATCATCCCCTTCTTCGTGAACCGCGTGGGC |
| CCCAAAAACGCTCTGCTGATCGGCGTGGTGATCATGGCTCTGCGCATCCTGAGCTGCGCTCTGTT |
| CGTGAACCCCTGGATCATCAGCCTGGTGAAACTGCTGCACGCTATCGAAGTGCCCCTGTGCGTGA |
| TCAGCGTGTTCAAATACAGCGTGGCTAACTTCGACAAACGCCTGAGCAGCACCATCTTCCTGATCG |
| GCTTCCAAATCGCTAGCAGCCTGGGCATCGTGCTGCTGAGCACCCCCACCGGCATCCTGTTCGAC |
| CACGCTGGCTACCAAACCGTGTTCTTCGCTATCAGCGGCATCGTGTGCCTGATGCTGCTGTTCGG |
| CATCTTCTTCCTGAGCAAAAAACGCGAACAAATCGTGATGGAAACCCCCGTGCCCAGCGCTATCTA |
| ACCATGCGAGAGTAGGGAACTGCCAGGCATCAAATAAAACGAAAGGCTCAGTCGAAAGACTGGGC |
| CTTTCGTTTTATCTGTTGTTTGTCGGTGAACGCTCTCCTGAGTAGGACAAAT |

**Table S4.** Oxidative stress indicators in free-growing cells and SS1 within ELM-SS1.

| Indicator | Free-growing cells | SS1 in ELM-SS1 |
|---|---|---|
| ROS (RFU/OD) | 17281.2 ± 2939.5 | 79178.0 ± 14834.0 |
| GSH (nmol/OD) | 2.293 ± 2.466 | 4.011 ± 2.301 |
| SOD (U/OD) | 192.35 ± 90.87 | 269.40 ± 63.27 |
| NADH (μM/OD) | 0.033 ± 0.002 | 0.044 ± 0.001 |
| $NAD^+$ (μM/OD) | 0.017 ± 0.005 | 0.013 ± 0.003 |
| $NADH/NAD^+$ | 2.02 ± 0.84 | 3.67 ± 1.05 |
| NADPH (μM/OD) | 0.07 ± 0.01 | 0.14 ± 0.02 |
| $NADP^+$ (μM/OD) | 0.15 ± 0.02 | 0.18 ± 0.04 |
| $NADPH/NADP^+$ | 0.48 ± 0.07 | 0.83 ± 0.24 |

**Table S5.** Representative GO terms in enrichment analysis of DEGs in ELM-SS1.

| GO ID | Term | p-value | Enrichment-score |
|---|---|---|---|
| **Upregulation** | | | |
| GO:0042597 | periplasmic space | 0.001715 | 4.240265 |
| GO:0022857 | transmembrane transporter activity | 0.056153 | 2.669797 |
| GO:0016730 | oxidoreductase activity, acting on iron-sulfur proteins as donors | 0.058201 | 4.805634 |
| GO:0009279 | cell outer membrane | 0.109507 | 3.432596 |
| GO:0009252 | peptidoglycan biosynthetic process | 0.132933 | 2.002347 |
| GO:0006979 | response to oxidative stress | 0.138291 | 3.003521 |
| GO:0071555 | cell wall organization | 0.174579 | 1.668623 |
| GO:0016491 | oxidoreductase activity | 0.217312 | 1.657115 |
| GO:0030288 | outer membrane-bounded periplasmic space | 0.231392 | 2.184379 |
| GO:0016020 | membrane | 0.588041 | 1.001174 |
| **Downregulation** | | | |
| GO:0006412 | translation | $5.73 \times 10^{-7}$ | 3.268199 |
| GO:0003735 | structural constituent of ribosome | $2.14 \times 10^{-7}$ | 3.582449 |
| GO:0042128 | nitrate assimilation | $1.27 \times 10^{-6}$ | 7.130617 |
| GO:0005840 | ribosome | 0.000149 | 3.034756 |
| GO:0030089 | phycobilisome | 0.000179 | 4.357599 |
| GO:0015934 | large ribosomal subunit | 0.002862 | 5.602627 |
| GO:0015979 | photosynthesis | 0.020336 | 1.782654 |
| GO:0015935 | small ribosomal subunit | 0.117922 | 3.268199 |
| GO:0043022 | ribosome binding | 0.117922 | 3.268199 |
| GO:0009772 | photosynthetic electron transport in photosystem II | 0.154373 | 2.801314 |

**Table S6.** Representative genes in KEGG analysis.

| Gene ID | Name | $Log_2$(Fold change) | p-value |
|---|---|---|---|
| **Oxidative stress** | | | |
| *RS08445* | *katG* | 3.022943 | $2.04 \times 10^{-49}$ |
| *RS11035* | *dpsA* | 1.682092 | $6.55 \times 10^{-13}$ |
| *RS02150* | *gst* | -1.00022 | $2.32 \times 10^{-6}$ |
| **Photosynthesis** | | | |
| *RS05140* | *psaD* | 1.000698 | $9.44 \times 10^{-20}$ |
| *RS06405* | *psaF* | 1.120881 | $2.02 \times 10^{-16}$ |
| *RS06400* | *psaJ* | 1.578382 | $6.14 \times 10^{-17}$ |
| *RS07110* | *psbA* | -2.46399 | $1.69 \times 10^{-81}$ |
| *RS03360* | *psbC* | -1.40967 | $6.85 \times 10^{-13}$ |
| *RS05330* | *psbP* | -1.10256 | $5.81 \times 10^{-5}$ |
| *RS05375* | *cpcA* | -1.46879 | $8.43 \times 10^{-10}$ |
| *RS05370* | *cpcB* | -1.68356 | $8.21 \times 10^{-12}$ |
| *RS05380* | *cpcC* | -1.64589 | 0.000362 |
| *RS05390* | *cpcE* | -2.03858 | $2.28 \times 10^{-14}$ |

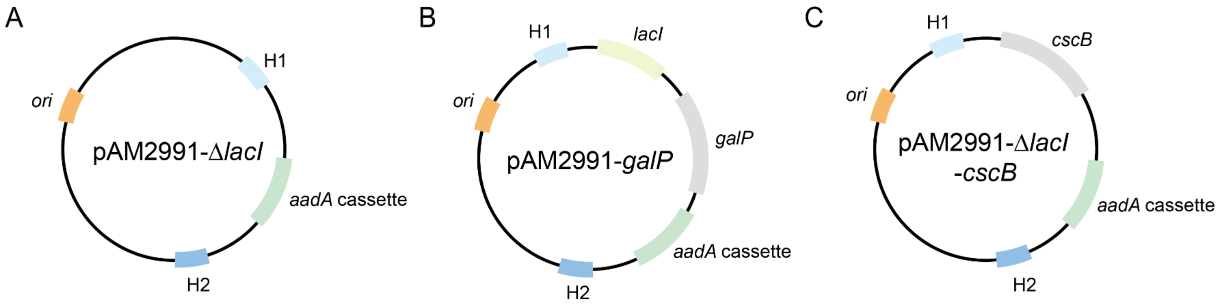


**Figure S1. Schematic diagrams of plasmids.** The plasmid pAM2991 was used for gene integration into the NS1 in the *S. elongatus* genome. **(A)** Plasmid pAM2991-Δ*lacI* was constructed via knocking out *lacI* cassette from pAM2991. **(B)** pAM2991-*galP* was constructed by inserting *galP* cassette into pAM2991, and *galP* was driven by the *lac*-$P_{trc}$ inducible system. **(C)** pAM2991-Δ*lacI*-*cscB* was generated by inserting *cscB* cassette into pAM2991-Δ*lacI*.

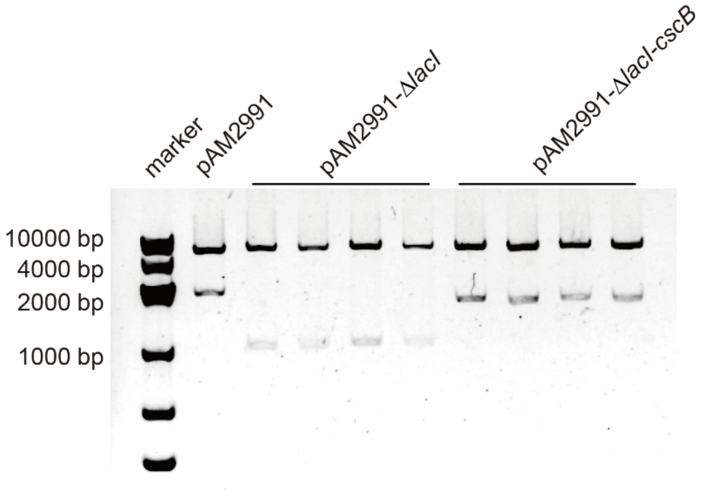


**Figure S2. Gel electrophoresis verification of plasmids pAM2991-Δ*lacI* and pAM2991-Δ*lacI*-*cscB*.** The plasmids were digested with Sac II at 30 °C for 1 h before verification.

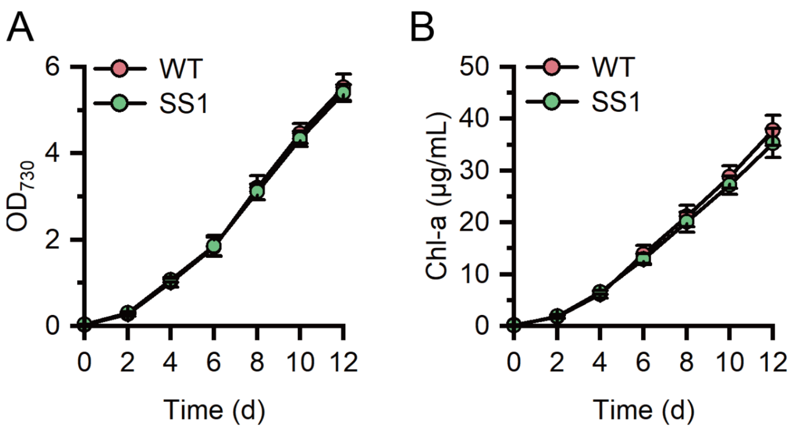


**Figure S3. Growth profiles of the wild-type strain and SS1. (A)** $OD_{730}$ and **(B)** the content of Chl-a of wild-type strain and SS1 during the 12 days of cultivation in BG11 medium. All experiments were performed in triplicate, and the error bars indicate the standard deviations of the means of biological replicates.

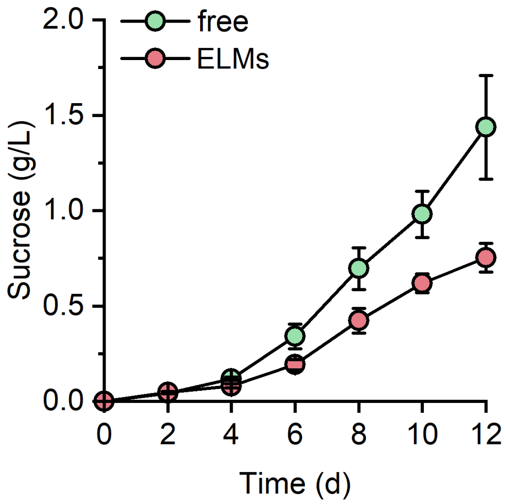


**Figure S4. Sucrose production of ELM-SS3 and the free-growing SS3.** The experiments were performed in triplicate, and the error bars indicate the standard deviations of the means of biological replicates.

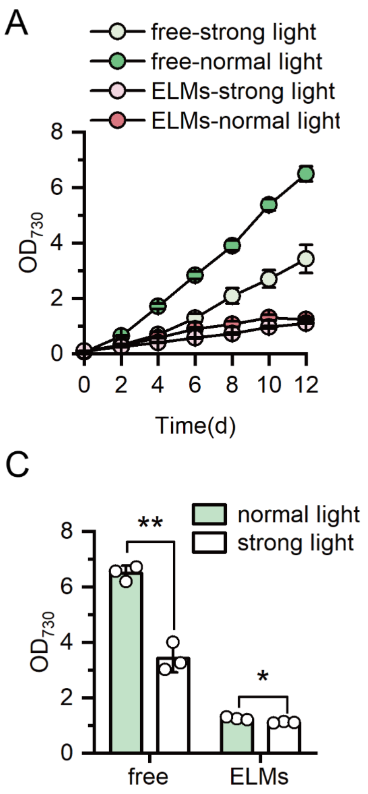


**Figure S5. Growth of SS2 within ELMs and planktonically. (A)** $OD_{730}$ and **(B)** the content of Chl-a of SS2 within ELM-SS2 and free-growing SS2 during the 12 days of cultivation at different light intensities. **(C)** $OD_{730}$ and **(D)** the content of Chl-a of SS2 in ELM-SS2 and growing freely on Day 12 at different light intensities, where 70 μmol photons·$m^{-2}$·$s^{-1}$ was regarded as high light intensity, and 35 μmol photons·$m^{-2}$·$s^{-1}$ was considered normal light intensity. The experiments were performed in triplicate, and the error bars indicate the standard deviations of the means of biological replicates. The differences were statistically assessed by the *t*-test (*$P$ < 0.05; **$P$ < 0.01).

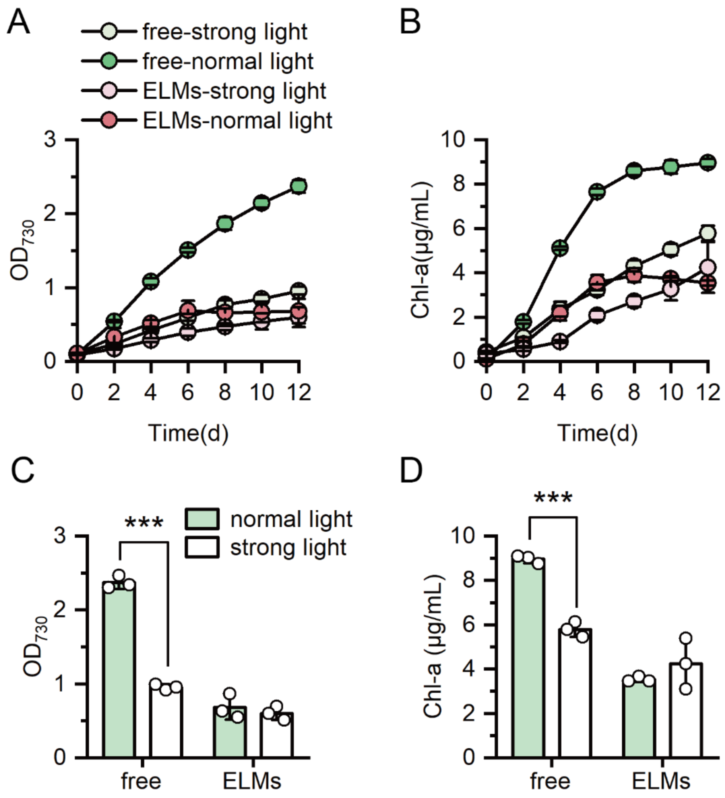


**Figure S6. Growth of SS3 within ELMs and planktonically. (A)** $OD_{730}$ and **(B)** the content of Chl-a of SS3 in ELM-SS3 and free-growing SS3 during the 12 days of cultivation at different light intensities. **(C)** $OD_{730}$ and **(D)** the content of Chl-a of SS3 within ELM-SS3 and growing freely on Day 12 at different light intensities, where 70 μmol photons·$m^{-2}$·$s^{-1}$ was regarded as high light intensity, and 35 μmol photons·$m^{-2}$·$s^{-1}$ was considered normal light intensity. The experiments were performed in triplicate, the error bars indicate the standard deviations of the means of biological replicates, and the differences were statistically determined by *t*-test (****P* < 0.001).

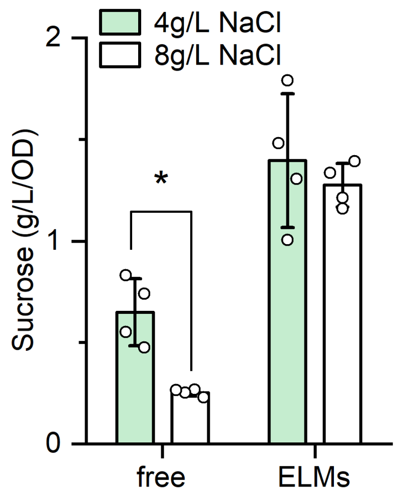


**Figure S7. The specific sucrose production of SS3 within ELM-SS3 and SS3 growing freely on Day 12.** The experiments were performed at least in triplicate, and the error bars indicate the standard deviations of the means of biological replicates, and the differences were determined by the *t*-test (*$P < 0.05$).

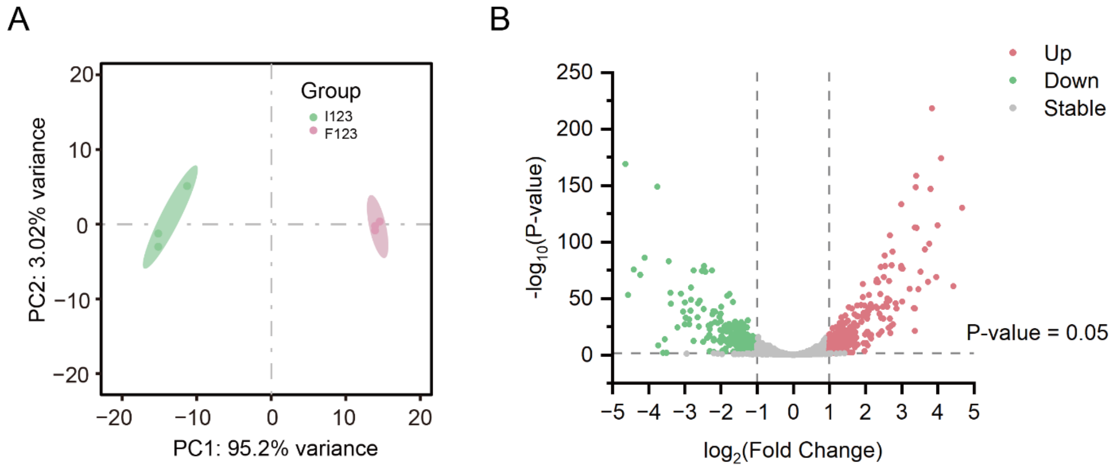


**Figure S8. Overall transcriptomic analysis. (A)** Principal component analysis (PCA) scatter plot of gene expression on SS1 in ELM-SS1 and free-growing cells, the green area represents the ELMs group and the red area represents the free-growing cells group. **(B)** Volcano maps of differentially expressed genes (DEGs) for the SS1 in ELM-SS1 and free-growing cells. Green dots represent the downregulated DEGs, red dots represent the upregulated DEGs, and gray dots represent the genes with no significant difference.

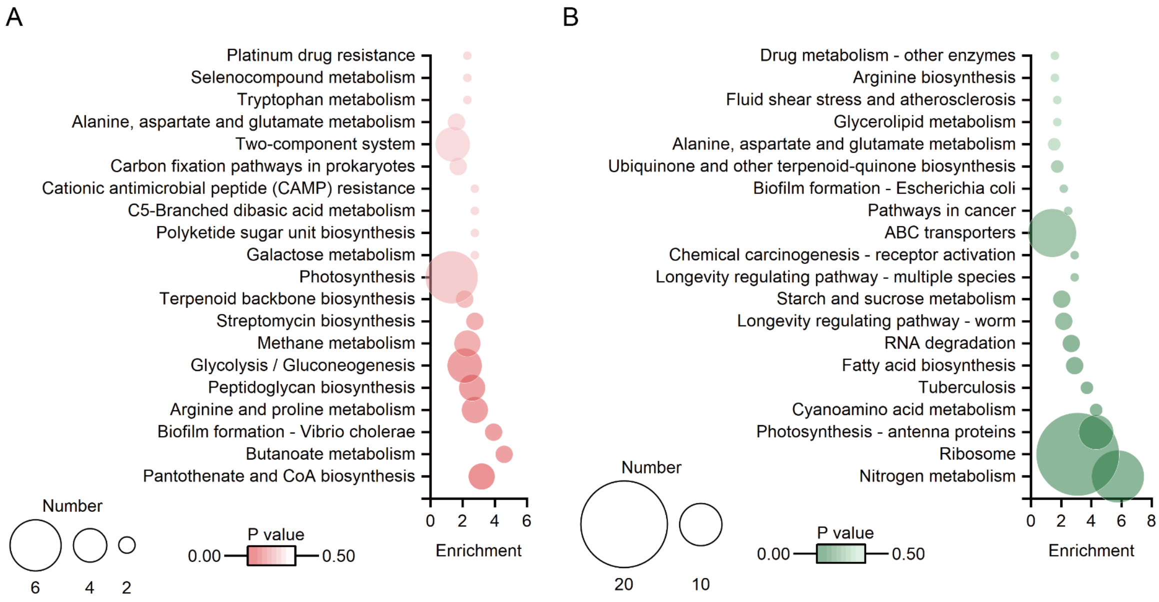


**Figure S9. KEGG enrichment analysis between SS1 within ELM-SS1 and growing freely. (A)** Representative upregulated KEGG of differentially expressed genes (DEGs). **(B)** Representative downregulated KEGG of DEGs. The color indicates P values, and the bubble size indicates the number of DEGs in KEGG.

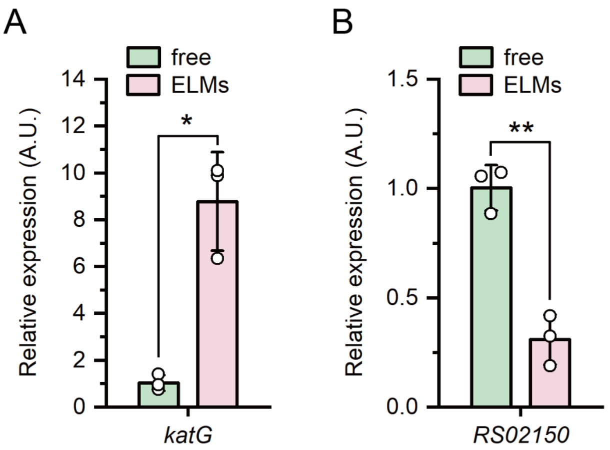


**Figure S10. Verification of RNA-sequencing data with qPCR. (A)** Relative expression of *katG* gene and **(B)** *RS02150* gene. The experiments were performed in triplicate, and the error bars indicate the standard deviations of the means of biological replicates. The differences were determined by the $t$-test (*$P < 0.05$; **$P < 0.01$).

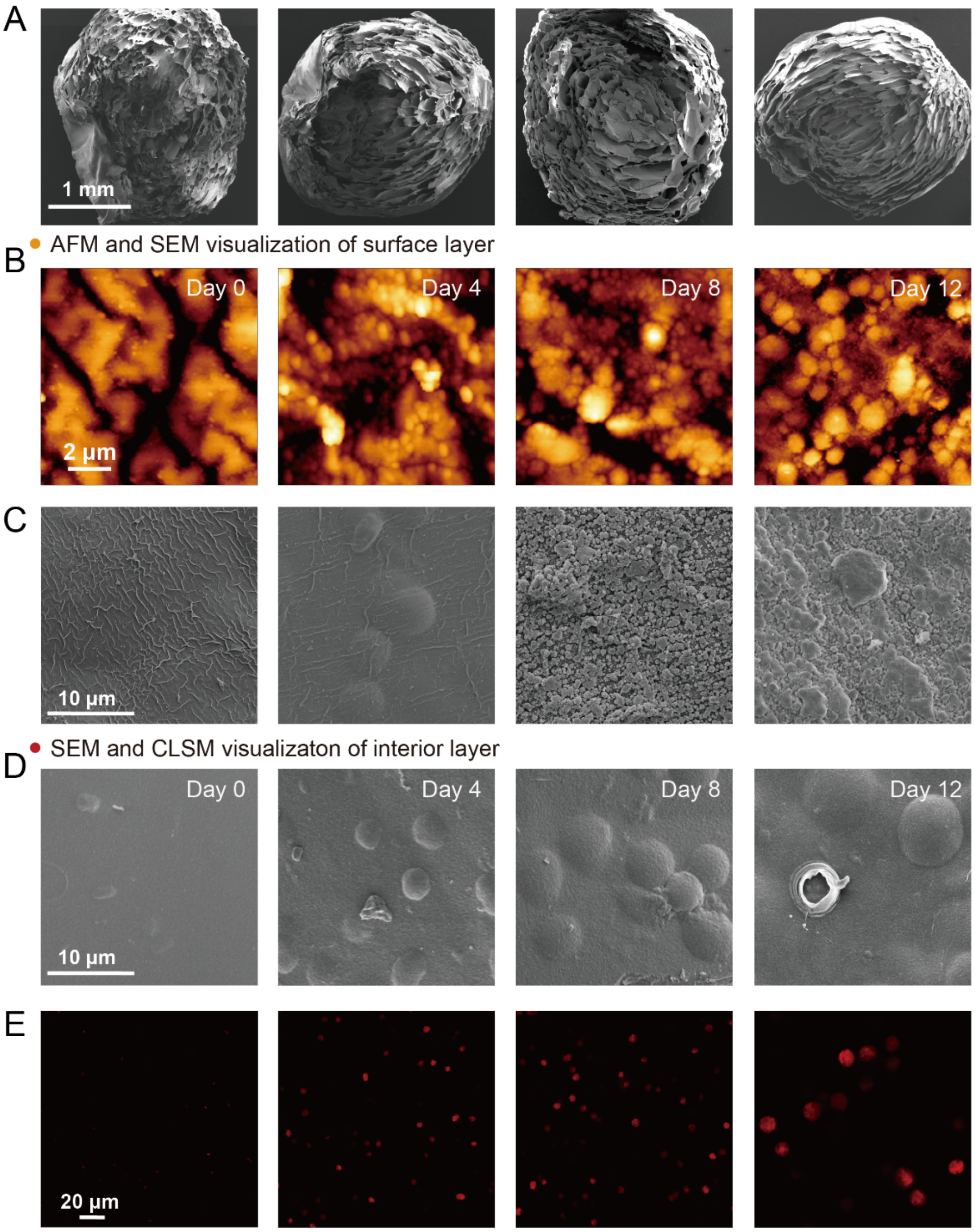


**Figure S11. Microscopical visualization of ELMs. (A)** The interior structure of ELM-SS1 visualized by SEM. **(B)** Surface morphology of ELM-SS1 visualized by AFM. **(C)** Surface morphology and **(D)** interior structure of ELM-SS1 visualized by SEM. **(E)** Representative CLSM images of ELM-SS1. Red fluorescence represents chlorophyll autofluorescence of *S. elongatus* cells. All analysis were performed on samples at 0, 4, 8 and 12 days of cultivation.
